# Common marmosets (*Callithrix jacchus*) do not differentiate between familiar and unfamiliar conspecifics based on pitch contour information

**DOI:** 10.1101/2025.10.27.684793

**Authors:** Julia Victoria Grabner, Emma Pigmans, Monika Mircheva, Job Morgan Knoester, Robert Baumgartner, Ulrich Pomper, Thomas Bugnyar, Michelle J Spierings

## Abstract

Previous studies suggest that vocal-learning species use variations of prosodic cues, such as changes in pitch, amplitude, or duration, in the conveying of meaning and emotions as well as individual recognition. Although it is known that the long-distance contact calls (“phee” calls) of common marmosets, a vocal non-learner, vary in prosodic information depending on individual characteristics, evidence of the species’ use of this information only recently started to emerge. In this study we tested 18 captive common marmosets’ (*Callithrix jacchus*) reactions to playbacks of familiar and unfamiliar individuals’ phee calls and extracted pitch-contour information. The playbacks consisted of i) natural phee calls from group members or unfamiliar conspecifics, or ii) synthesized pitch contours of familiar or unfamiliar phee calls following a natural or artificial syllable order as well as pure tones of similar frequency and duration. We found that although individuals seemed to show different reactions towards the natural calls of familiar and unfamiliar individuals’, we did not find group level differences. Marmosets furthermore show significantly reduced interest in synthesized pitch contours compared to natural phee calls and did not differentiate between pitch contour playbacks and pure tone control stimuli, indicating that they did not categorize the pitch contours as conspecific calls. Our results suggest that extracted pitch contour information alone is not sufficient for marmosets to recognise conspecifics, however further studies are required to investigate the exact mechanism of individual recognition in marmosets.

## Introduction

While living in social groups offers advantages like predator defence and foraging success (e.g., Rubenstein, 1978), it also introduces complex cognitive challenges. Hence, social interactions within and outside of a group have been proposed as one of the main drivers of cognitive evolution (Ashton et al., 2020; Byrne & Whiten, 1988; Dunbar, 1998). One cognitive challenge is to individually identify conspecifics. Differentiating between group members and outsiders, as well as individually recognising them and remembering their previous interactions with them, is crucial to navigate social life, cooperate and compete with others. Indeed, studies in various social primate species found that individuals can differentiate between group members, familiar outgroup individuals (i.e. neighbouring individuals) and unfamiliar individuals and adjust their reactions accordingly, showing for example differences in aggression or retreat behaviour between neighbours and strangers (e.g., French et al., 1995; Herbinger et al., 2009; Wich et al., 2002). However, the exact characteristics that are used to differentiate between individuals vary between species and situations and often remain unclear.

Depending on their environment and species-specific traits, individuals may rely on visual (e.g., Parr et al., 2000; Pokorny & de Waal, 2009a; Pokorny & de Waal, 2009b), olfactory (e.g., Henkel & Setchell, 2018; Smith et al., 1997) or acoustic information (e.g., Cheney & Seyfarth, 1980; Herbinger et al., 2009; Pereira, 1986; Rendall et al., 1996) or a combination of those modalities to discriminate between individuals. Acoustic signals seem especially important in species living in dense habitat, big complex groups or groups spread over large territories (Fichtel & Manser, 2010; Xie et al., 2024). Not only the vocalisation type itself, but also prosodic cues – variations in syllable duration, pitch and amplitude – prove to be important to provide context (see for example Filippi (2016) for a review on emotional prosody). Studies show that animals adapt their calling rates and acoustic properties of their vocalisations according to their surroundings (e.g., Ey et al., 2009; Koda et al., 2008; Sugiura, 2007). Furthermore, acoustic signals allow the extraction of several characteristics of an animal, such as their emotional state (Briefer, 2012), approximate size, sex, age and dominance status (e.g., Ey et al., 2007; Fischer et al., 2004; Norcross et al., 1999) as well as individual identity (e.g., Marler & Hobbett, 1975). Despite these individual differences in vocalisations and the fact that several animal species can distinguish individuals based on their vocalisations, studies investigating which characteristics individuals use are sparse.

One study on greater false vampire bats (*Megaderma lyra*) using synthesized conspecific calls showed that they could differentiate calls with different prosodic cues and mainly used pitch to do so (Janßen & Schmidt, 2009). Further evidence about the importance of different prosodic cues comes from human and animal studies using discrimination tasks based on human speech syllables investigating which information is used to distinguish between acoustic signals. While humans can distinguish between different made-up words using only amplitude or pitch alone, Budgerigars (*Melopsittacus undulatus*) require more than one prosodic cue to distinguish two different stimuli (Hoeschele & Fitch, 2016). In a similar study Zebra finches (*Taeniopygia guttata*) performed comparably to humans, being able to use a single prosodic cue for differentiation between syllables, with pitch being the most salient cue (Spierings & ten Cate, 2014). While it seems that vocal-learning species display advanced abilities, rats (*Rattus norvegicus*), a vocal non-learning species, were shown to only be able to discriminate between human speech syllables when pitch, duration, amplitude and vowel quality information were available (Toro & Hoeschele, 2017). Just removing one of those prosodic cues led to very poor performance, suggesting a potential link between vocal learning and the perceptive abilities of prosodic cues. However, whether the same prosodic cues are also important in intraspecific communication and whether a similar difference between vocal learners and vocal non-learners exists remains unknown.

Common marmosets’ (*Callithrix jacchus*) long-distance contact calls (“phee” calls) exhibit differences in prosodic-like features that are context- and sex dependent (Norcross & Newman, 1993) as well as group- and individual-specific (Miller et al., 2010; Zürcher & Burkart, 2017; Zürcher et al., 2019). Playback studies furthermore show that marmosets differentiate calls from different conspecifics (Kato et al., 2018; Miller & Thomas, 2012; *Callithrix jacchus*) and seem to categorise calls depending on callers’ sex (Norcross et al., 1994; *Callithrix jacchus*) or familiarity (Rukstalis & French, 2005; *Callithrix kuhlii*). This suggests that vocalisations encode characteristics that allow individuals to individually recognize conspecifics. Although it has been found that common marmosets’ vocalisations show individual characteristics in prosodic information and that individuals can differentiate between conspecifics, it is not known if they are able to differentiate conspecifics from a single salient cue or if they, as a vocal non-learner, need the complete prosodic information.

In this study, we used a playback paradigm to investigate whether common marmosets use prosodic information to individually identify conspecifics. We first investigated if marmosets show a difference in vocal response and looking time towards i) familiar and unfamiliar conspecifics’ natural unaltered contact (“phee”) calls, ii) extracted pitch information of those phee calls, as well as iii) control sounds with similar acoustic properties. As common marmosets are territorial and show strong reactions towards unfamiliar individual’s phee calls (e.g., Caselli et al., 2018) we predicted that individuals would show clear differences in looking times with longer looking times during playbacks of unfamiliar phee calls compared to familiar calls and controls. Furthermore, as marmosets are known to engage in vocal turn-taking (Takahashi et al., 2013), we predicted that individuals would vocally respond to the playbacks. We expected them to vocalize more often after familiar individuals’ calls, compared to unfamiliar individuals’ calls and pure tone control stimuli. For the extracted pitch information, we had two alternative expectations: if pitch is the prominent information used by individuals, we expected individuals to show similar patterns of responses when presented with unaltered playbacks or extracted pitch information. However, if animals need more acoustic information to distinguish conspecifics, we would expect individuals to not show any differentiation between playbacks of familiar and unfamiliar individuals’ pitch contours.

## Material and Methods

### Subjects

For this study we tested 18 adult common marmosets (6 f/12 m; mean ± SD: 9.4 ± 4.9 years) at the Animal Care Facility of the Faculty of Life Sciences, University of Vienna, Vienna, Austria. Individuals were housed in family groups of three to seven individuals in three separate keeping rooms. Enclosures were equipped with various items such as branches, wooden boards, (sleeping) baskets, hammocks, foraging boxes, heating- and UV-lamps and had bark mulch floor covering. Additionally, individuals had access to roofed outdoor enclosures when temperatures allowed. Enclosures were regularly enriched with a variety of food- and non-food items. The monkeys were fed three times a day with a rich selection of vegetables, animal protein and fruits and had ad libitum access to water. Gummi arabicum as well as additional vitamin- and mineral supplements were provided daily.

### Experimental procedure and stimuli

We performed a playback experiment during which each tested marmoset was presented with a set of stimuli including altered and unaltered vocalisations of familiar and unfamiliar conspecifics. We then measured the reaction of the tested individual towards those stimuli by scoring behavioural responses. The experiment was divided into two conditions which were tested with the same individuals in consecutive weeks: the Natural call condition, and the Pitch contour condition, that are explained in more detail below. Before, between and after the stimuli pink noise of random durations (10-30 seconds) was played to avoid habituation. Familiar stimuli were vocalisations from individuals of the same family group. Unfamiliar stimuli were vocalisations from individuals from separate colonies in Zurich and Tübingen which the test subjects have never heard or seen before. All vocalisation stimuli were recorded during habituation sessions while a single individual was separated from its family group. Recordings were edited in Praat (version 6.2.14; Boersma, 2001) to match in amplitude, and single stimuli were cut and combined into one playback. In each test session the calls of a different familiar and unfamiliar conspecific were presented, while within a test sessions different calls of the same familiar and unfamiliar individual were used. Example stimuli can be accessed on our Open Science Framework repository (https://osf.io/kn3pt).

### Natural call condition

In the Natural call condition, we wanted to find out how individuals react to playbacks of natural calls from familiar and unfamiliar individuals. To do this, recordings of phee calls of familiar and unfamiliar conspecifics as well as control sounds were used. Playbacks consisted of natural three-syllable loud phee calls or three pure tones with similar properties (1 second pure tones, 7 – 8.4 kHz, 10 ms ramp-up/ramp-down, 300ms silence) as control (see figure 1*a,b* for spectrograms of example stimuli). Each test session contained four randomized sets of stimuli, each set comprising one familiar individual’s phee call, one unfamiliar individual’s phee call and one control sound, making a total of 12 stimuli per session.

**Fig. 1.**
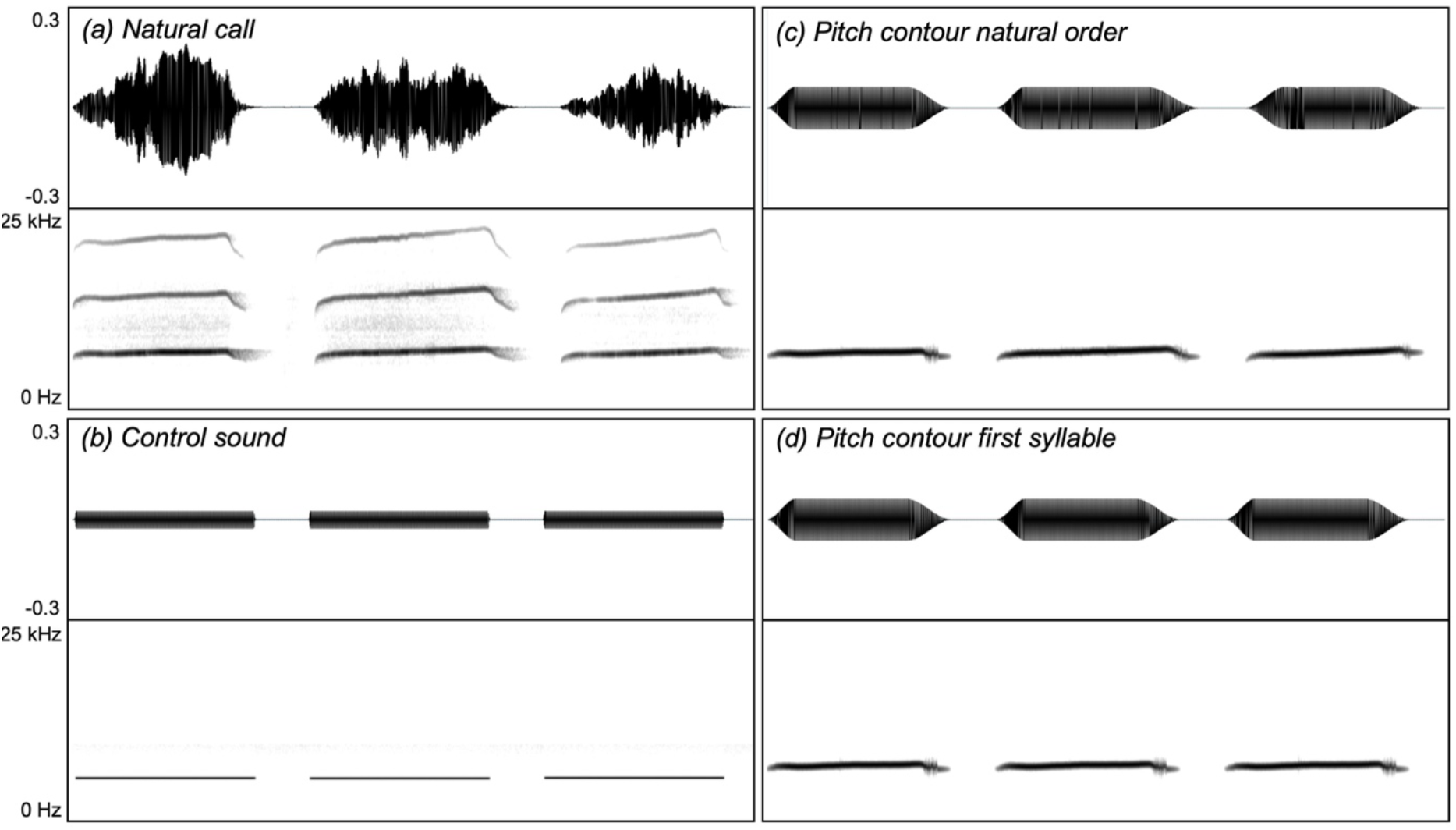
Types of playback stimuli used in the study. (a) natural call composed of three phee call syllables in their natural order, (b) control sound composed of three pure tones of similar duration and frequency, (c) pitch contour natural order, composed of pitch contours of the three syllables of a phee call in the natural order, (d) pitch contour first syllable, composed of three times the pitch contour of the first syllable of a phee call.

### Pitch contour condition

In the Pitch contour condition, we investigated marmosets’ reactions to extracted pitch information of natural phee calls. Here, the pitch contour of each syllable of the call was extracted and saved separately. This was done using the built-in Praat function “extract pitch tier”. The extracted pitch contour, containing only the fundamental frequency, was then synthesized to sound (sine). After resynthesizing, the single syllables were concatenated to a three-syllable string with 300ms of silence between syllables. In this step two different stimulus types were produced: the *natural order* stimuli, that consisted of all three syllables of the natural call in original order (1-2-3), and the *first syllable only* stimuli, in which only the first syllable was used and repeated three times to exclude further temporal information (1-1-1) (see figure 1*c,d* for spectrograms of example stimuli). In each session of the Pitch contour condition three randomized sets, each containing one familiar natural order pitch contour, one familiar first syllable only pitch contour, one unfamiliar natural order pitch contour, one unfamiliar first syllable only pitch contour and one control sound were presented, resulting in a total of 15 stimuli per session.

### Experimental setup and data collection

In both conditions subjects were tested individually in a separate research room. After entering the cage compartment (1m x 1m x 1m) through a tunnel system, the individual sat on a small perch facing a loudspeaker. The experimenter then hid behind a curtain and started the trial. The stimuli were played back from a loudspeaker (Anchor Explorer Pro) placed 1.5m from the experiment compartment and hidden behind a fake plant, using Audacity (version 3.2.5) on a MacBook pro13. Volume levels on Audacity, the laptop and the speaker were kept constant throughout the experiment, such that the sound had a mean sound pressure level of 50dB in the middle of the experimental cage. All experimental sessions were filmed using two cameras: The main camera (Canon legria HF G25) was placed on top of the speaker directly facing the experimental cage to measure the time the individuals look in the direction of the sound source. Camera 2 (Canon legria HF R806) was mounted above the cage for a top view (figure 2). Recordings from camera 2 were only used when an individual was out of view from the main camera. Fourteen individuals were tested in three test sessions for each condition (Natural call and Pitch contour). Each individual was tested once a week over the course of 8 weeks (three weeks data collection for Natural call condition, two weeks break, three weeks data collection for Pitch contour condition). Due to changes in group composition four individuals were only tested in one (Lokri) or two (Kobold, Luna and Jack) test sessions per condition over the course of 4 or 6 weeks, respectively.

**Fig. 2.**
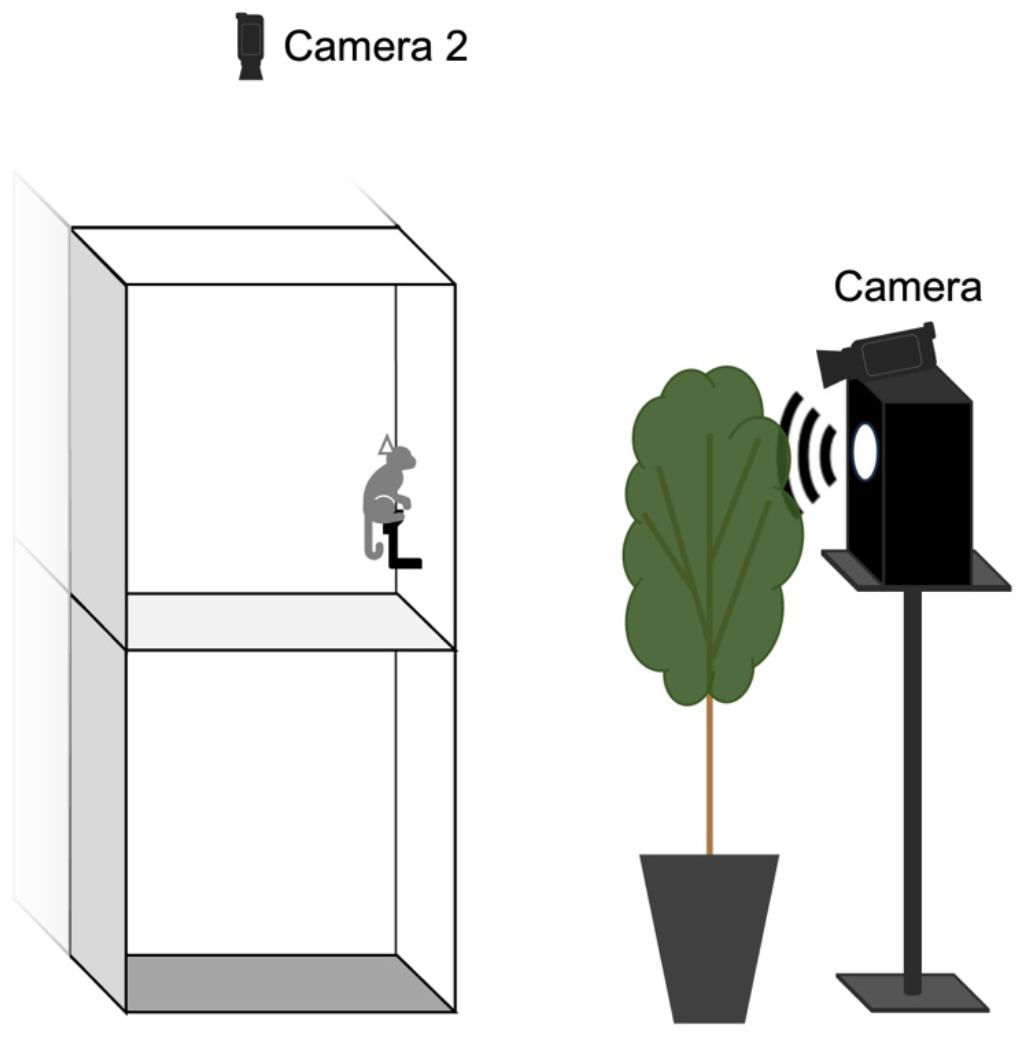
Schematic of the experimental setup. The stimuli were played back from a speaker hidden behind an artificial plant. On top of the loudspeaker the main camera was placed to record the focal individual’s reactions. Over the experimental cage an additional camera (camera 2) was placed.

### Video coding

The videos were analysed by a single observer (E.P.) in BORIS (version 7.13; Friard & Gamba, 2016). The observer coded each video twice. In the first round all videos were analysed without sound to code looking time, the amount of time the individual looked in the direction of the speaker, while being blind to the stimulus category and onset. In the second round, the frequency of contact (“phee”) call vocalisations and stimuli on- and offset were coded. To measure vocal responses to the playback we coded all phee calls occurring between the end of the stimulus and 10 seconds after. Due to different lengths of stimuli and individuals sometimes being out of sight we used proportions of looking times for the analysis. 10% of the videos were additionally coded by a second observer (J.V.G.) to ensure interobserver reliability (IOR). Interobserver reliability was excellent for overall duration (two-way intraclass correlation coefficient: ICC (A,1) = 0.998) and frequency (ICC(A,1) = 0.972) of coded variables, as well as looking time durations (ICC(A,1) = 0.921). Reliability was good for frequency of phee call vocalisations (ICC(A,1) = 0.897).

### Statistical analysis

#### General approach

Data extraction and structuring as well as statistical analysis were done in R (version 4.5.0; R Core Team, 2025) using RStudio (version 2024.12.1+563; Posit team, 2025). As response variables we used the proportion of time the individuals were looking at the speaker (looking time proportion) as well as a binary response (yes/no) variable indicating whether the individuals responded by looking at the speaker (looking response) or producing phee calls within ten seconds of the stimulus (vocal response). For the analysis we used Generalised Linear Mixed Models (GLMM) with the function glmmTMB (Brooks et al., 2017) and applied the full-null model comparison framework, where we compared a full model including all test predictors, representing the working hypothesis against a null model excluding all test predictors, representing the null hypothesis, to control for cryptic multiple testing (Forstmeier & Schielzeth, 2011). If the model comparison was significant, meaning the sum of test predictors significantly improved the model fit, we continued with model diagnostics and investigated the model results. If the model comparison was not significant, we halted analysis and did not further investigate the model. If the model comparison showed trend-level effects (*p* values over 0.05 but below 0.1) we decided to follow the approach proposed by Titchener et al. (2023), to not ignore the possibility of an effect and report both the full model and the reduced model as well as the resulting analysis. We do however advice the readers to treat these results with caution.

Before the model comparison, we performed model diagnostics. Firstly, we evaluated the variance inflation factors (VIFs) using the function vif (of the package “car”, version 3.1-3) to investigate the extent of collinearity of predictors. Secondly, we checked model dispersion using the custom function overdisp.test (Mundry, 2024 personal communication). For binominal models we additionally produced basic diagnostic plots using the package DHARMa (version 0.4.7; Hartig, 2024). Lastly, we visually inspected the plots of the best linear unbiased predictors (BLUPs). These were also used to aid in the decision which random slopes to first exclude in the case of model convergence problems. If model diagnostics showed acceptable model fit the model results were investigated using the summary() function to compare levels within a predictor against the reference. The drop1() function (with the argument test = “Chisq”) was then used to assess the influence of each fixed effect by excluding one fixed effect at a time and comparing this reduced model to the full model including the effect. Where applicable pairwise post hoc tests were conducted using the emmeans package (version 1.11.0; Lenth, 2025) and adjusted using Tukey correction. As an additional step we investigated model stability by dropping one level of a specified grouping factor (in our case “subject”) and comparing the resulting estimates against each other using the function influence_ mixed from the package glmmTMB.

#### Looking response / - time

First, we investigated differences in looking time proportions between stimuli within the Natural call condition as well as the Pitch contour condition. Because of substantial zero inflation of the response variable looking time proportion, we opted for a two-step approach: First, we transformed the response variable to the binominal variable looking response, representing a binary yes/no response. We then ran a GLMM with a binominal distribution to compare between reaction and no reaction. Then, we excluded zeros and ran an additional GLMM with beta regression, to test whether looking time proportion differed between stimuli.

For both the binominal and the beta model, we included as fixed effects test predictors the interactions of stimulus type with the subject’s sex, and stimulus type with the subject’s age including their main effects. We included the session number and set number as fixed effects as control predictors. As random effects we included individual identity, family group, session number and set number, plus all theoretically identifiable random slopes using the function fe.re.tab (Mundry, 2023). The resulting null models included the same fixed and random effects control predictors, but not the fixed effects test predictors. In case the following full-null model comparison was significant, but an interaction had no significant effect, we formulated a null model lacking that interaction, but containing its main effects, to check for an effect of those main effects in absence of an interaction. In the case of model convergence issues, we simplified the model stepwise: First we removed correlations between random effects intercepts and random slopes. Then we excluded random slopes based on their biological, ecological relevance and their expected importance in the experimental design. Once a full model converged and we had to further simplify the random slopes because of convergence issues in the null model or drop1 sub-models, we also used the BLUPs to assist in our decision making.

### Vocal response

To investigate differences in vocal responses between stimuli within the Natural call condition and the Pitch contour condition we again first transformed the response variable to the binominal variable vocal response, representing a binary yes/no response. We then used a GLMM with a binominal distribution including the same fixed and random effects as for looking responses. As multiple phee call vocalisations were nearly never observed (only 4 out of 588 cases in the Natural call condition and only 3 out of 735 cases in the Pitch contour condition) we did not run further analyses for the frequency of phee call vocalisations.

### Comparison Natural call condition and Pitch contour condition

To compare looking responses and -times between the Natural call condition and Pitch contour condition we again used a GLMM with binominal and a GLMM with beta regression, both including the experimental condition as test predictor and including the same fixed and random effects as for looking responses but not including interactions with the subject’s sex or age as we did not expect any interactive effects between the experimental condition and the individuals’ characteristics.

## Results

### Natural call condition

#### Common marmosets show interest in conspecific’s natural phee calls

In the Natural call condition, for the binominal model investigating looking responses (responses versus no responses) the full model including stimulus performed significantly better than the null model (χ^2^(6) = 16.02, *p* = 0.014). Looking responses differed significantly between the stimulus categories (χ^2^(2) = 13.09, *p* = 0.001) (figure 3*a*). A Tukey-adjusted post-hoc pairwise comparison revealed that individuals responded more often when a conspecific call was played back compared to a control stimulus (control vs familiar: estimate = -0.99, *SE* = 0.24, *z* = -4.14, *p* < 0.001; control vs unfamiliar: estimate = -1.00, *SE* = 0.34, *z* = -2.90, *p* = 0.011). However, although density plots indicate more interest in unfamiliar calls over familiar ones (figure 4a) we did not find a difference in looking responses between the familiar and unfamiliar stimulus (estimate = -0.01, *SE* = 0.34, *z* = -0.02, *p* = 0.999). For the beta model investigating looking time proportions, the full model including stimulus type did not perform significantly better compared to the null model (χ^2^(6) = 1.41, *p* = 0.965).

**Fig. 3.**
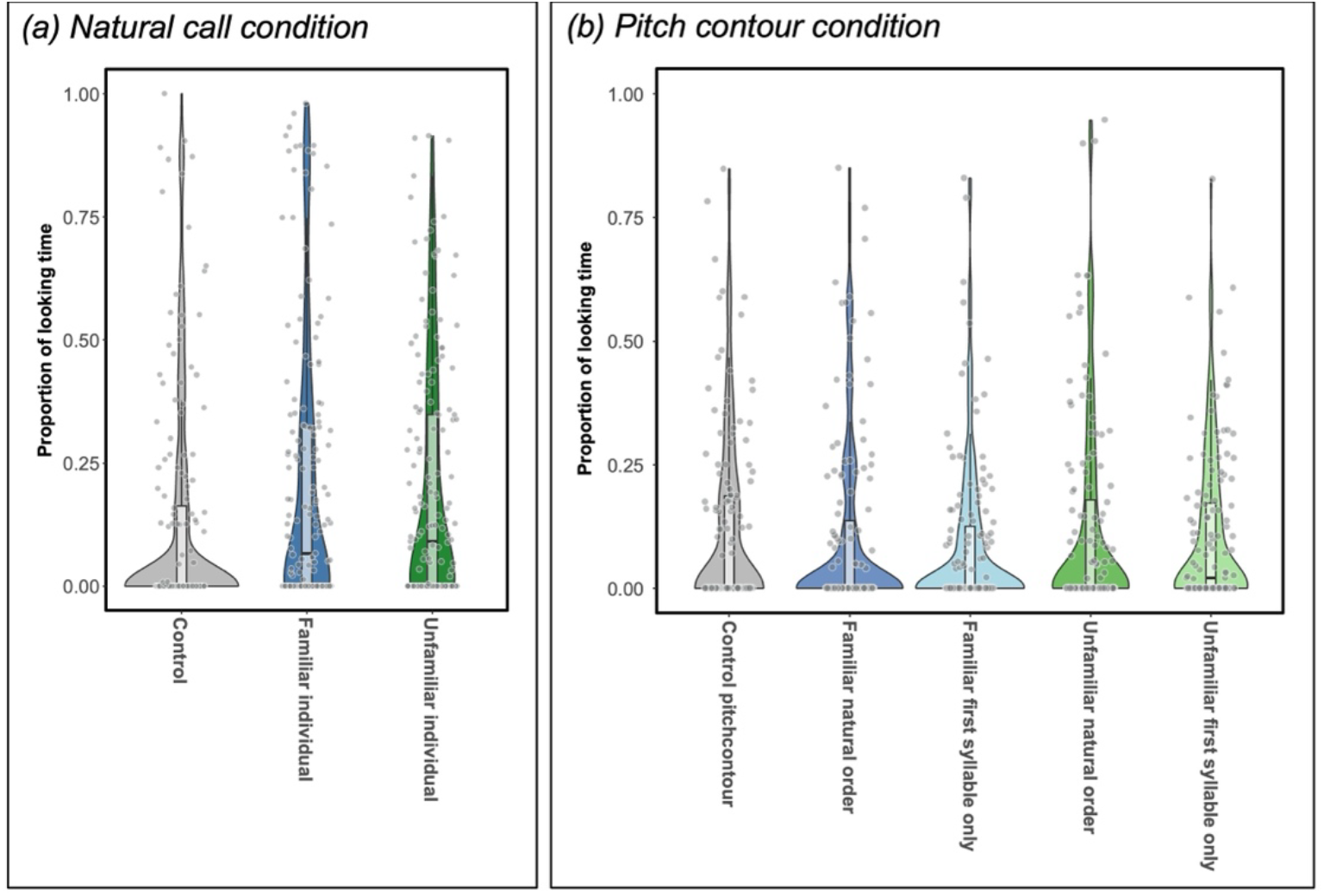
Proportion of time looking at the playback location per stimulus and experimental condition. (a) Natural call condition, (b) Pitch contour condition. While in the Natural call condition significant differences between familiar and unfamiliar individual’s calls compared to the control sound can be observed, looking time proportions did not differ between stimuli in the Pitch contour condition.

**Fig. 4.**
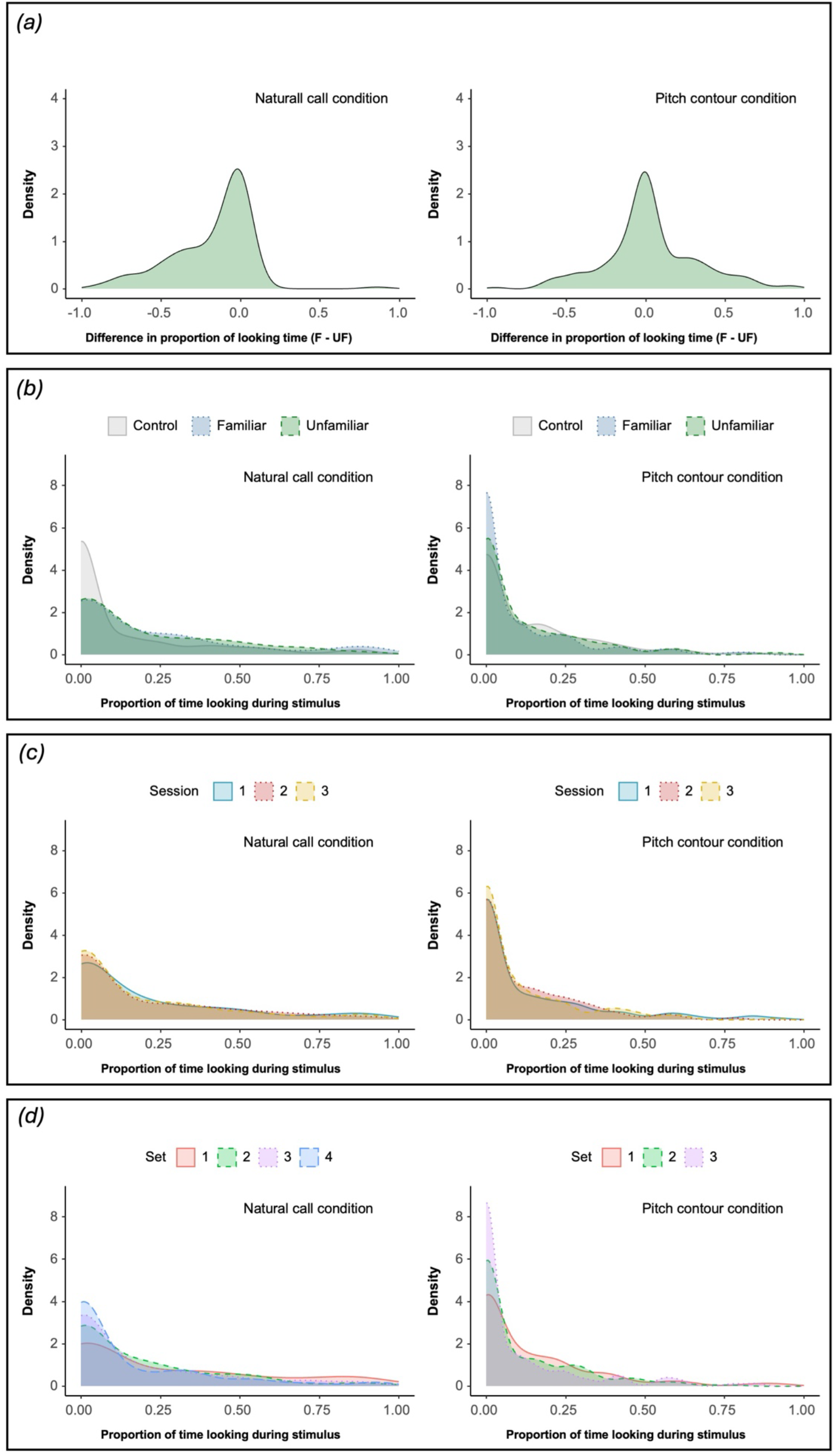
Density plots for the difference in proportion of looking between familiar and unfamiliar individuals and the total proportion of looking time. (a) shows the overall difference in the proportion of looking times between familiar and unfamiliar stimuli. The plot of the Natural call condition shows a more skewed distribution indicating a slight preference in looking at unfamiliar stimuli, while the plot in the Pitch contour condition is more uniformly distributed. (b), (c) and (d) show the proportion of time looking during the stimulus presentations over stimuli category (b), testing session (c) and set number (d) respectively. All three show higher densities around 0 in the Pitch contour condition, indicating less interest in the stimuli in the Pitch contour condition compared to the Natural call condition.

### Common marmosets react differently to familiar and unfamiliar individual’s natural phee calls on an individual level

Looking at figure 5a, it seems that although on a group level no difference in reaction between familiar and unfamiliar calls was found, on an individual level marmosets might react differently to familiar and unfamiliar conspecifics’ phee calls, but in different directions and to different extents. To investigate a possible effect of individual characteristics on the looking response and -times we included the individuals’ age and sex in the analysis. We did not find a significant effect of those characteristics on overall looking responses in the binominal model (sex: χ^2^(1) = 0.001, *p* = 0.978; age: χ^2^(1) = 0.04, *p* = 0.836). Interactions between sex or age and the stimulus did not significantly improve the binominal model fit (stimulus:sex: χ^2^(2) = 2.11, *p* = 0.348; stimulus:age: χ^2^(2) = 0.22., *p* = 0.897), and graphical inspection does not show any clear pattern (figure 6).

**Fig. 5.**
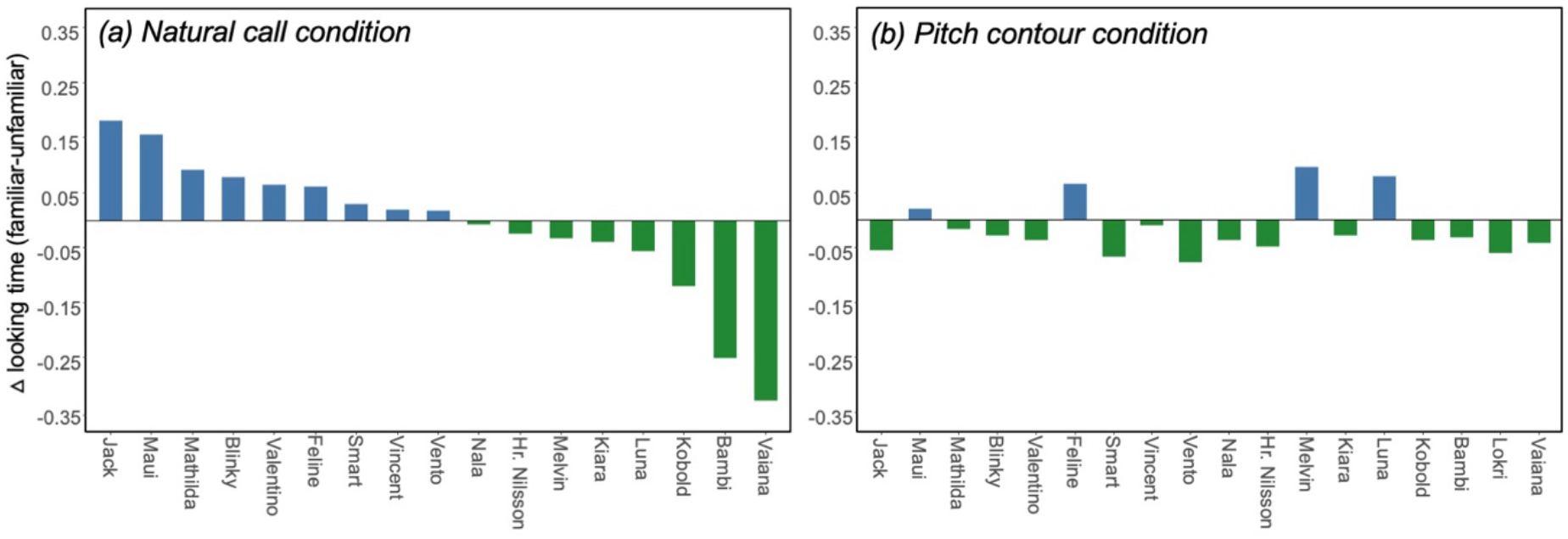
Delta values comparing times looking at the playback location between familiar and unfamiliar stimuli per individual (familiar – unfamiliar). In blue (above 0) individuals that looked longer at familiar stimuli, in green (below 0) individuals that looked longer at unfamiliar stimuli. (a) Natural call condition, ordered by degree of interest from most distinctly interested in familiar stimuli to most distinctly interested in unfamiliar stimuli. (b) Pitch contour condition, in the same order as the Natural call condition. In the Pitch contour condition overall decreased differences between familiar and unfamiliar stimuli can be observed. Individuals furthermore don’t show the same looking patterns as in the Natural call condition.

**Fig. 6.**
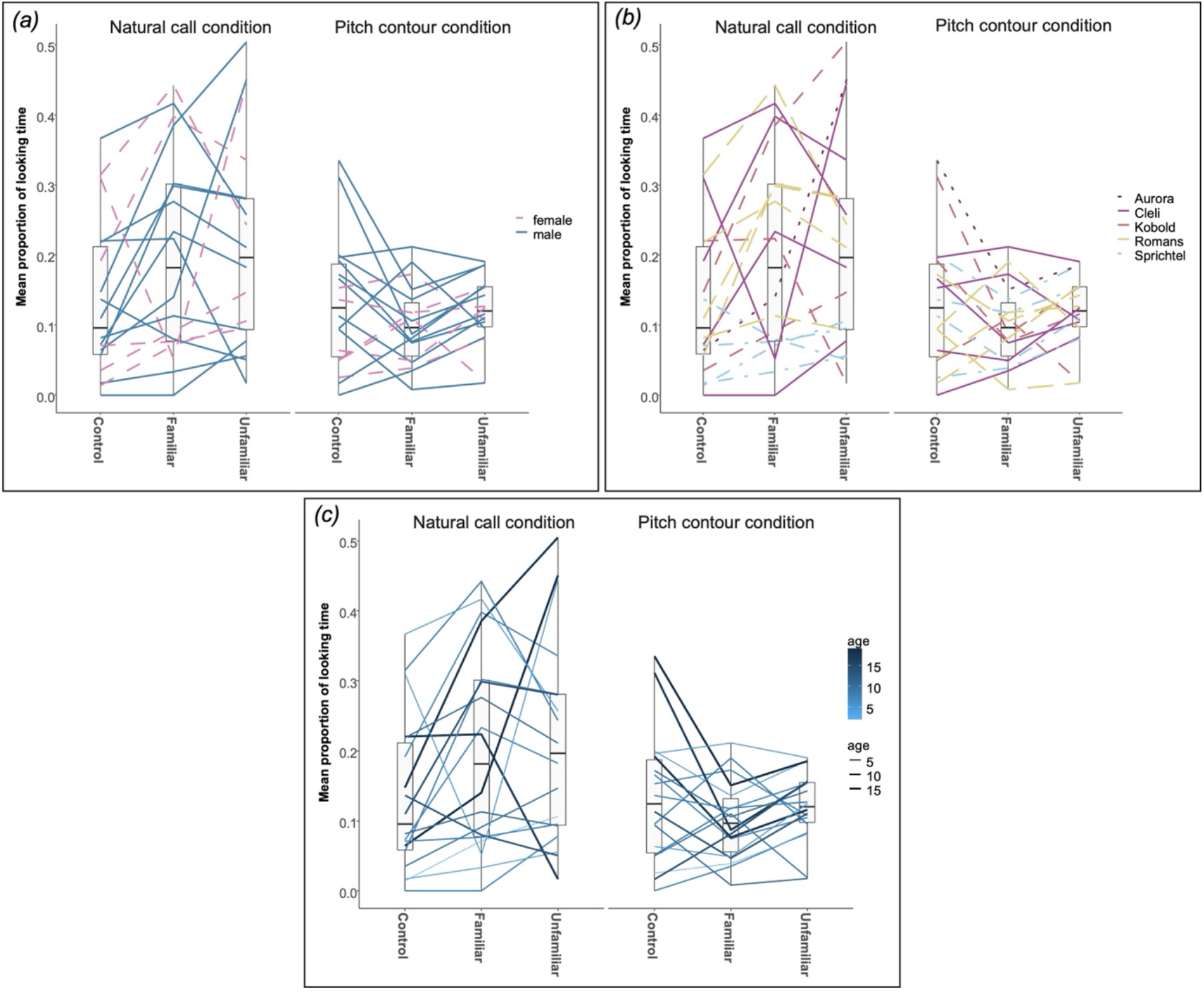
Effects of (a) sex, (b) family group and (c) age on the mean proportion of looking time per stimulus (control, familiar, unfamiliar), both in Natural call and Pitch contour condition. No clear patterns could be found for either of the variables.

### Marmosets rarely vocally respond to familiar and unfamiliar individual’s playbacks

Overall vocal responses were rare, with individuals only producing phee calls in response to 138 of 588 total stimuli. For the binominal model investigating the effect of vocal responses the full model including the stimulus type did not perform significantly better compared to the null model (χ^2^(6) = 8.11, *p* = 0.230).

### Pitch contour condition

#### Common marmosets do not react differently to familiar and unfamiliar individual’s pitch contours on a group level

In the Pitch contour condition, the full binominal model including stimulus did not improve the model fit compared to the null model for both looking response (χ^2^(12) = 6.90, *p* = 0.864) and vocal response (χ^2^(12) = 8.61, *p* = 0.736). When excluding zero responses, the full beta model for looking time proportions again did not perform significantly better than the null model (χ^2^(12) = 19.25, *p* = 0.083). However, as the model comparison showed a trend level effect, we report the model results but advice the readers to carefully evaluate these findings. As interactions did not significantly improve the model fit (stimulus:sex: χ^2^(4) = 4.11, *p* = 0.391; stimulus:age: χ^2^(4) = 3.99, *p* = 0.408) we fitted a reduced model including only the main effects without interactions. We found that looking time proportions significantly differed between stimuli (χ^2^(4) = 9.81, *p* = 0.044). However, post-hoc pairwise comparisons did not yield significant differences (figure 3b).

#### Common marmosets do not react differently to familiar and unfamiliar individual’s pitch contours on an individual level

We could not observe clear individual preferences in looking time proportions in the Pitch contour condition. Differences in reactions between familiar and unfamiliar stimuli seem to be considerably less pronounced compared to the Natural call condition and preferences do not match the ones observed in the Natural call condition (figure 5b). Similarly to the Natural call condition, sex (χ^2^(1) = 0.09, *p* = 0.769) and age (χ^2^(1) = 0.23, *p* = 0.632) did not significantly influence overall looking time proportions (figure 6).

### Comparison Natural call condition and Pitch contour condition

#### Common marmosets show less interest in pitch contours compared to natural call playbacks

The observed lack of differentiation between stimuli in the Pitch contour condition could be attributed to a loss of interest or habituation effect. Indeed, for both the binominal model investigating looking responses and the beta model investigating looking time proportions the full model including the experimental condition performed significantly better than the null model (looking responses: χ^2^(1) = 7.27, *p* = 0.007; looking time proportions: χ^2^(1) = 23.68, *p* < 0.001). Looking responses (χ^2^(1) = 7.27, *p* = 0.007) and looking time proportions (χ^2^(1) = 23.68, *p* < 0.001) both significantly decreased between the Natural call condition and the Pitch contour condition (figure 4b). However, while individuals’ interest decreased between testing sets within a session (looking response: χ^2^(1) = 5.23, *p* = 0.022; looking time proportion: χ^2^(1) = 3.72, *p* = 0.054) there was no significant decrease in looking over testing sessions (looking response: χ^2^(1) = 0.02, *p* = 0.902; looking time proportion: χ^2^(1) = 1.03, *p* = 0.311), indicating only short time habituation or loss of interest (figure 4c,d). Furthermore, we found a significant increase in looking time proportions (χ^2^(1) = 4.08, *p* = 0.043) but not looking responses (χ^2^(1) = 0.02, *p* = 0.876) with age.

## Discussion

We found that individuals’ reactions towards familiar and unfamiliar stimuli did not differ when presented with pitch contour information only. This suggests that pitch information alone is not sufficient for marmosets to differentiate between conspecifics. Interestingly however, they also did not show different reactions between conspecific pitch contours and control stimuli. The overall decrease of looking time proportions between the Natural call condition and the Pitch contour condition suggests that marmosets were not interested in the pitch contour stimuli.

### The reduced interest suggests an inability to perceive pitch contours as conspecific calls

As we observed a considerable decrease in attention between the two conditions but no significant decrease between testing sessions within each condition, it is unlikely, that the decrease is due to a habitation effect. Rather it seems that the pitch contours were not perceived as conspecific calls, leading to individual’s interest dropping to control stimulus levels. One possibility is that the manipulation to extract the pitch contours made them too artificial for the individuals to be interesting. However, studies show that marmosets react similarly to natural and synthesized phee calls (e.g., Norcross et al., 1994). On the other hand literature suggests, that while vocal learners can distinguish between sounds using only limited prosodic information (Hoeschele & Fitch, 2016; Janßen & Schmidt, 2009; Spierings & ten Cate, 2014), vocal non-learners need the full spectrum of prosodic cues for the same task (Toro & Hoeschele, 2017). If the same is true for intraspecific communication, excluding prosodic information would make it impossible for vocal non-learning individuals such as marmosets to distinguish between calls and could possibly lead to the observed loss of interest in the Pitch contour condition. While the current study design does not allow to disentangle between a possible effect of artificial sound manipulation and the general ability to extract information from one prosodic cue alone, we show that the marmosets clearly respond less to phee calls when only the fundamental frequency is maintained.

### Individual characteristics are likely to lead to differing responses

Surprisingly, in contrast to previous studies investigating marmoset’s reactions to conspecific calls (e.g. *Callithrix jacchus*: Miller & Wang, 2006; *Callithrix kuhlii*: Rukstalis & French, 2005) we did not find a categorical difference in reaction towards familiar and unfamiliar individuals’ calls. However, our data indicates there might be various effects of individual differences. Based on the graphical representation it seems that in the Natural call condition most individuals preferably react to one of the two categories. This preference however seems to vary in direction and strength between individuals which might lead to no clear pattern emerging on the group level. The discrepancy in reaction compared to what Rukstalis and French (2005) found could be caused by the fact that in our study additional individual characteristics could play a role. While they only used playbacks of pair bond partners or unfamiliar individuals of the opposing sex, we used familiar and unfamiliar playbacks originating from both males and females of varying age (2-17 years) and each reproductive status (breeders, helpers and pair-kept individuals without breeding experience). Furthermore, relationships between the individuals and playbacks used in this study varied. Individuals heard familiar calls from their pair partners, siblings, parents or offspring. According to previous studies individuals’ reactions depend both on their own characteristics as well as the characteristics of the played back individual and an interplay of both. For example, Caselli et al. (2018) found that, when being played back unfamiliar individuals’ calls, breeding individuals reacted first to playbacks of the opposite sex and breeding females reacted with pilo-erection to playbacks of unfamiliar females. Another study from Norcross and colleagues (1994) also shows differences in reaction, with breeding individuals reacting significantly stronger than juveniles and differences in the exhibited behaviours between males and females. Although our sample size and the number of potentially influencing variables did not allow us to investigate demographic effects in more detail, the possible presence of individual preferences might suggest that marmosets were able to differentiate between familiar and unfamiliar individual’s calls.

### The absence of vocal responses highlights the importance of playback structure

Another surprising result was the lack of vocal responses. The tested individuals rarely vocalized within the set time window of 10 seconds after the playbacks. Although, to our knowledge, no study directly investigated differences in vocal response behaviour towards familiar and unfamiliar playbacks, we expected individuals to strongly respond with phee calls to conspecific vocalisations as shown in previous studies. In a study by Norcross and colleagues (1994) marmosets produced twice as many phee calls after than before the playback of an unfamiliar individual’s phee call, but did not show significant differences in response rates towards playbacks from different sexes or call variants. The way the playback was presented in our study might have influenced the individual’s responses. Previous studies found, that individuals do not react differently to playbacks of phee calls and silent playbacks when the playback is not presented in an interactive way (Miller & Wang, 2006) and when latencies between an individuals’ phee call and playbacks are too long (Miller et al., 2009). Furthermore, it was found, that individuals responded less often after the identity of the played back individual changed, than when the identity of the counterpart stayed the same (Miller & Thomas, 2012). In our study each playback session consisted of calls from two different individuals played back in a semi-random order. Moreover, playbacks were not presented in an interactive way but in a fixed timescale and order, resulting in individuals sometimes starting to vocalize shortly before the start of a stimulus and leading to overlaps between the vocalisations and the playbacks. The comparison between our findings and the literature shows that the way playbacks are presented can have a strong influence on the exposed subjects’ response.

### Age is unlikely to drive the observed effects

The overall low number of observed reactions raises the question whether the individuals perceived the stimuli correctly. One potential factor influencing the marmosets’ reactions could be their age. Marmosets above 8 years of age are often classified as “aged” and exhibit diverse symptoms of age-related pathologies (Abbott et al., 2003; Tardif et al., 2011). Studies show that older age is related to hearing-loss in marmosets (Harada et al., 1999; Sun et al., 2021). Two reasons however speak against this explanation: Firstly, age-related hearing loss in marmosets is described to be only moderate (Harada et al., 1999) and primarily affects ultra-high frequencies above 16kHz (Sun et al., 2021). It is therefore unlikely to influence the perception of phee calls that have a substantially lower fundamental frequency (5.5-10.5 kHz; Norcross & Newman, 1993). Secondly, if hearing loss would be the reason for the lack in response, we would expect individuals to also show overall declined reactions and no differentiation in reaction towards control stimuli and conspecific stimuli. We do however find that overall proportional looking time increased with age and that individuals do differentiate between control and conspecific stimuli in the Natural call condition. Taken together, although age might be an influencing factor it cannot fully explain the observed results. Rather, although we cannot statistically show it, we think the effect might stem from individuals displaying individual differences in their reactions and them not classifying the pitch contours as conspecific calls.

### Our results give limited insight into individual recognition in marmosets

Our results only partially shed light on the acoustic characteristics used in individual recognition in marmosets. While it is known from previous studies that marmosets can differentiate between conspecifics based on vocalisations alone (Kato et al., 2018; Miller & Thomas, 2012) we set out to investigate which acoustic cues are used to do so. Marmosets are vocal non-learners and one vocal non-learning species has been shown to perform worse when differentiating sounds based on only one prosodic cue at least when using human speech syllables (Toro & Hoeschele, 2017). However, it is not clear yet whether vocal learning and the ability to use certain acoustic cues is connected. We therefore expected them to either show similar or less pronounced differences in reactions towards familiar and unfamiliar conspecifics’ calls, depending on whether the inability to use single prosodic cues expands to marmosets. In line with the latter predictions, we found that marmosets do not react differently towards extracted pitch contours of familiar and unfamiliar individuals. However, given that the individuals did not show clear differentiation between familiar and unfamiliar calls in the Natural call condition and did not differentiate between pitch contours and control sounds we cannot conclude whether marmosets need more information or even the full acoustic spectrum or whether the observed behaviour results from other factors such as habituation, loss of interest or too artificial stimuli.

## Acknowledgements

We thank Steffen Hage, University of Tübingen for providing phee call recordings used for the unfamiliar stimuli. Furthermore, we are grateful to Awani Bapat, Christian Blum and David Meijer for valuable assistance in data analysis.

## Conclusion

Taken together our results suggest that marmosets react differently to playbacks of natural familiar and unfamiliar conspecifics’ contact calls on an individual basis. On a group level though, overall effects are outweighed by these individual preferences. When being presented with only the extracted pitch contours however, individuals seem to not differentiate anymore. Additionally, individuals showed no differentiation between extracted pitch contours and pure tone control sounds. Further studies are needed to disentangle whether marmosets cannot use pitch alone to distinguish between individuals or whether simply extracting the pitch contour and playing back a resynthesized version is too artificial for the monkeys to be perceived as a conspecific call.

